# “Biophysical and Colloidal Properties Govern Anti-Drug Antibody Generation: A Case Study Using the Tumoricidal mAb 4713”

**DOI:** 10.1101/2025.09.03.674127

**Authors:** Sanjida Yesmin, M. Monirul Islam, Md. Din Islam, Hiromichi Tsurui, Tomonori Saotome, Shuji Matsuoka, Yutaka Kuroda

## Abstract

Anti-drug antibodies (ADA) are a well-recognized clinical problem, but their sporadic occurrence has limited detailed biophysical characterization of causative factors. We investigated how different stress conditions influence aggregation of a murine monoclonal antibody, mAb 4713, and the resulting potential to induce ADA in mice. Dynamic light scattering (DLS) revealed that stirring, freeze–thaw, and heat stress each generated subvisible aggregates (30–700 nm), whereas non-stressed mAb 4713 remained predominantly monomeric. Stirring- and freeze–thaw–induced aggregates retained native-like secondary and tertiary structures, thermal and biochemical stability, and anti-tumor cytotoxicity, as shown by circular dichroism, tryptophan fluorescence, thermal shift assay, limited proteolysis, and cell death assay. In contrast, heat-induced aggregates displayed complete structural denaturation and loss of cytotoxic activity. Immunization of Jcl: ICR mice demonstrated that heat-induced aggregates were only mildly immunogenic, while freeze–thawed and non-stressed mAb 4713 elicited no detectable immune response, and ADA generated under these conditions did not affect anti-tumor activity. Strikingly, stirred aggregates triggered a strong anti-mAb antibody response, as measured by ELISA, and the resulting antisera completely abolished mAb 4713 cytotoxicity against Raji cells, a Burkitt lymphoma cell line, indicating induction of neutralizing ADA. These results show that aggregate immunogenicity depends not simply on aggregate presence but critically on their structural and colloidal properties and formation pathways, highlighting the need to assess both structural and colloidal properties of aggregates in biopharmaceutical development to mitigate ADA risks.

## 1. Introduction

Protein therapeutics are becoming increasingly important for the treatment of life-threatening diseases, including cancer and autoimmune disorders [1]. Although amino acid sequences are usually optimized to minimize immune responses against foreign proteins, aggregation can still trigger strong immunogenicity, potentially compromising therapeutic efficacy [2–6]. This unwanted immune response is characterized by the generation of anti-drug antibodies (ADA) against the therapeutic protein drugs. Neutralizing ADAs have been shown to impact therapeutic proteins by diminishing or entirely blocking their activity, whereas non-neutralizing ADAs do not have such effect [4,5,7] and are thus not harmful, *per se*. Nevertheless, the sporadic nature of ADA induction has resulted in limited experimental data in animal models, hindering a comprehensive understanding of the phenomenon [3,4,8].

Protein aggregation is a major factor in inducing unwanted immune responses and subsequent ADA generation, and they may arise from minor perturbations at any stage, from protein production to patient administration [8–11]. These perturbations include slight temperature variations, changes in solvent conditions, or mechanical stresses, all of which can disrupt protein structure and stability or lead to aggregate formation [10,12–14]. Furthermore, the strength and nature of the applied stresses influence the properties of the resulting aggregates, including their size (soluble oligomers, subvisible, and visible aggregates), conformation (native-like, partially unfolded, or fully unfolded), reversibility (reversible or irreversible), colloidal stability, and morphology [15]. Several studies have examined the role of aggregate size, conformation, and other biophysical properties in ADA induction both *in vivo* and *in vitro* [16–20]. However, the extent to which the colloidal and biophysical properties of protein aggregates contribute to ADA formation, specifically the formation of neutralizing ADA, which lead to the loss of therapeutic activity, rather than the mere production of non-neutralizing antibodies remains poorly understood.

Monoclonal antibodies (mAbs) have emerged as a leading class of protein therapeutics due to their high specificity and strong binding affinity for the target molecules [9,21]. As aforementioned, small perturbations can promote aggregation of mAbs, trigger immune responses, and lead to ADA formation, ultimately compromising their therapeutic activity [4,7]. Here, we sought to characterize the biophysical, colloidal, and biochemical properties of the aggregates generated under well-controlled conditions by applying artificial stresses (heating, stirring, and freeze-thawing) that mimic those encountered by mAbs during their life cycle.

To this end, we used a murine-derived anti-pan HLA class II monoclonal antibody, mAb 4713, as a model system. The mAb 4713, whose humanized form is currently in advanced development as an anti-cancer therapeutic, has been extensively characterized [22]. We generated distinct aggregate types of mAb 4713 and assessed their role in triggering unwanted immune responses, focusing particularly on the induction of neutralizing ADA as measured by the loss of anti-tumor cytotoxicity. We observed that native-like, colloidally stable aggregates elicited a stronger immune response than the less stable freeze-thawed and heat denatured aggregates in a mouse model. Furthermore, the antisera produced by the stirred aggregates, neutralized the therapeutic ability of mAb 4713 to kill malignant lymphoma cells, which demonstrated the production of ADA. These findings suggest that the conformational and colloidal properties of mAb aggregates can significantly influence neutralizing ADA production and impair the therapeutic efficacy of mAbs, underscoring the need for further investigation in this area.

## 2. Materials and Methods

### 2.1. Preparation of mAb 4713 aggregates

The mAb 4713 was established by Matsuoka et al. [22] and Burkitt lymphoma cell line (Raji cell) was obtained from Riken BRC Cell Bank (Tsukuba, Japan). We applied three different stresses, heating, stirring and freeze-thawing to produce aggregates using 1.5 mL centrifuge tubes at a concentration of 0.3 mg/mL in 1x phosphate-buffered saline (PBS), pH 7.4, as previously reported [19]. Precisely, the protein samples were separately heated at 70 °C for 30 minutes, stirred using magnetic stirrer bar at 200 rpm for 30 minutes at 25 °C, and frozen once using liquid nitrogen for 3 minutes and thawed at 25 °C to generate heat (Ht), stirred (St), and freeze-thawed (Ft) aggregates, respectively. Non-stressed mAb 4713 (Ns) was used as a control in all experiments.

### 2.2. Biophysical measurements

We assessed the biophysical, thermal, and biochemical characteristics of the non-stressed monomers and stressed aggregates of mAb 4713 under the same conditions of 0.3 mg/mL protein concentration in 1x PBS, pH 7.4, as used in immunological studies.

The dynamic light scattering (DLS) measurements were conducted using a Malvern Zetasizer Nano-S system (Malvern, UK) and a polystyrene cuvette at 25 and 37 °C. The hydrodynamic radius (*R*_h_) of the mAb 4713 aggregates was calculated from the number distributions utilizing the Stokes-Einstein equation [23] and averaged over three readings.

The static light scattering (SLS) intensity was measured using an FP-8500 spectrofluorometer (JASCO, Tokyo, Japan) and a 3 mm path-length cuvette, exciting the samples at 600 nm, followed by recording emission in the wavelength range of 550-650 nm at 25 and 37 °C. Each sample was measured thrice, and the values were averaged.

We evaluated the secondary structures by far-ultraviolet (UV) circular dichroism (CD) spectroscopy using a JASCO J820 CD spectropolarimeter (JASCO, Tokyo, Japan). The CD spectra were measured with a continuous scanning wavelength ranging from 205 to 260 nm, using a 2 mm path-length quartz cuvette (TOSOH, Tokyo, Japan) at 25 and 37 °C.

We investigated the tertiary structure and molten globule characteristics by tryptophan and 8-anilino-1-naphthalene-sulfonic acid (ANS) spectroscopy, using a 3 mm path-length cuvette and an FP-8500 spectrofluorometer (JASCO, Tokyo, Japan). The tryptophan fluorescence spectra were measured at 25 and 37 °C with an excitation at 295 nm, and the emission spectra were recorded from 300 to 500 nm.

For ANS fluorescence, ANS (Sigma-Aldrich, Steinheim, Germany) was first added to the sample at a final concentration of 20 μM and then incubated for 20 minutes in the dark at 25 °C. Finally, the spectra were recorded with the emission over the wavelength of 400-600 nm, with an excitation wavelength set at 380 nm at 25 and 37 °C.

We assessed the thermal stability by thermal shift assay (TSA) using qTOWER3G (Analytikjena, Endress+Hauser, Germany). Precisely, the samples were prepared by adding 2.5 µL of 8000x diluted Sypro orange dye to 17.5 µL of non-stressed monomers and stressed aggregates, according to manufacturer protocols. Next, the temperature-dependent thermal shift (melting curve) was monitored in a temperature range of 25-80 °C with a heating rate of +0.1 °C/s [24]. Finally, the melting temperature was calculated using qPCRsoft 4.1 software.

To investigate thermal stability and structural integrity of non-stressed monomers and stressed aggregates, the differential scanning calorimetry (DSC) samples were prepared as described previously [25] and dialyzed for 18 hours at 4 °C in 1x PBS (pH 7.4) using a Spectra/Por 3 membrane (MWCO of 3.5 kDa) with one buffer exchange. After dialysis, the protein concentrations of samples were adjusted to 0.3 mg/mL, followed by degassing of the samples. DSC measurements were performed using a VP-DSC microcalorimeter (MicroCal, Malvern Panalytical, Malvern, UK) at a scan rate of +1.0 °C/min in the temperature range of 20-100 °C. Blank measurements were taken using 1x PBS buffer, pH 7.4 before measurements.

We evaluated the biochemical stability through limited proteolysis using trypsin (Nacalai Tesque Inc., Kyoto, Japan) as reported previously [19]. Shortly, the non-stressed monomers and stressed aggregates at a concentration of 0.15 mg/mL were treated with trypsin at a final concentration of 175 µg/mL and incubated at 37 °C. Sampling was done at different time points and analysed by SDS-PAGE. The protein fractions in the bands were calculated from the SDS-PAGE image by CS analyzer 4 software (ATTO, Tokyo, Japan).

### 2.3. Mice immunization studies

We evaluated the immunogenicity of the non-stressed monomers and stressed aggregates of the mAb 4713 in the Jcl: ICR (CLEA, Tokyo, Japan) mouse model. A total of 12 female mice of 4 weeks of age were divided into four groups (Supplementary Table S1). All mice groups were injected subcutaneously three times at two-week intervals at 30 µg of mAb 4713/dose/mouse in 1x PBS, pH 7.4. Blood samples were collected through tail-bleeding (TB) at weekly intervals (Supplementary Fig. S1).

On day 50, mice were anesthetized with isoflurane (a small piece of cotton soaked with 300 μL of isoflurane in a 1L beaker for 1-2 minutes), and heart blood was collected via cardiac puncture using a 22G needle. Following blood collection, the mice were euthanized through cervical dislocation, and the spleens were harvested for flow cytometry analysis.

### 2.4. ELISA

We assessed the anti-mAb 4713 antibody (ADA) responses by enzyme-linked immunosorbent assay (ELISA), as previously reported [19,26–28]. The 96-well microtiter plates (Thermo Fisher Scientific, Roskilde, Denmark) were coated with monomeric non-stressed mAb 4713 at 100 ng/mL (IgM), 40 ng/mL (IgG) and 10 ng/mL (IgG1, IgG2a) concentrations in PBS, following overnight incubation at 25 °C. After washing once with PBS, wells were blocked using 1% bovine serum albumin (BSA; Wako, Osaka, Japan) in PBS for 45 minutes at 37 °C. Dose-specific mouse antisera were applied at an initial dilution of 1:125 (IgM) and 1: 250 (IgG), 1: 500 (IgG1, IgG2a) followed by a 3-fold serial dilution in 0.1% BSA-PBS and incubated for 2 hours at 37 °C. The plates were washed thrice with PBS-0.05% Tween-20 and once with PBS. Anti-mouse antibody-HRP conjugates (Thermo Fisher Scientific, Illinois, USA) were added at a dilution of 1: 2,000 (IgM, IgG2a), 1: 3,000 (IgG), and 1: 25,000 (IgG1), according to manufacturer protocols, in 0.1% BSA-PBS-Tween-20 and incubated for 90 minutes at 37 °C. Finally, 100 µL of ortho-phenyl-di-amine substrate (0.4 mg/mL in 50 mM citrate-phosphate buffer, pH 5.0 with 4 mM hydrogen peroxide) was added. The reaction was stopped with 1 N sulfuric acid after 20-minute incubation and colour intensity was measured at 492 nm (OD_492nm_) using a microplate reader (SH-9000 Lab, Hitachi High-Tech Science, Tokyo, Japan). Anti-mAb 4713 antibody titers were calculated using a power-fitting of OD_492nm_ versus the reciprocal of the sera dilution using a cutoff of OD_492nm_ = 0.1 above the background values and averaged over all mice in each group. The ELISA data were analysed using MS Excel (Microsoft, Redmond, WA, USA).

### 2.5. Flow cytometry analysis

The expression of cell surface cluster of differentiation (CD) markers (CD44 and CD62L) on and intracellular cytokines (IFN-γ and IL-4) expression of mouse splenic T-lymphocytes (helper T cells, T_H_, and cytotoxic T cells, T_C_) was analysed by flow cytometry, as previously reported [19,29]. In brief, first, a single-cell suspension of mouse splenocytes collected on day 50 was prepared in fluorescence-activated cell sorter (FACS) buffer (PBS supplemented with 2% fetal bovine serum (FBS), and 1 mM EDTA), and red blood cells (RBC) were lysed using RBC lysis buffer (0.15 M ammonium chloride, 10 mM potassium bicarbonate, and 0.1 mM EDTA). Lymphocytes were then labelled for T_H_ and T_C_ lineages separately (T_H_ lineage: anti-CD3-PC5.5, anti-CD4-PC7, anti-CD44-FITC, and anti-CD62L-PE; T_C_ lineage: anti-CD3-PC5.5, anti-CD8-PC7, anti-CD44-FITC, and anti-CD62L-PE) following the manufacturer protocols (BioLegend, San Diego, CA, USA).

Similar protocol was followed for intracellular cytokines labelling but in the presence of 0.05% Tween-20 (T_H_ lineage: anti-CD3-PC5.5, antiCD4-PC7, and anti-IFN-γ-PE/IL-4-PE) [29]. After a 30-minute incubation in the dark, unbound conjugated antibodies were removed by washing with FACS buffer and resuspended in FACS buffer. Finally, flow cytometry analysis was done using a CytoFlex (Beckman Coulter, Indianapolis, USA).

### 2.6. Cell death assay

We investigated the anti-tumor cytotoxicity i. e., the therapeutic activity of mAb 4713 using Raji cells. The interaction of mAb 4713 with its target receptor, HLA class-II present on Raji cells, ultimately results in cell death, which was assessed through flow cytometry (Beckman Coulter, Indianapolis, USA) and by counting cells under a light microscope (Nikon ECLIPSE Ts2-FL, Tokyo, Japan) as previously reported [22]. Raji cells were grown in RPMI-1640 media (Wako, Osaka, Japan) supplemented with 10% FBS at 37 °C and 5% CO_2_ up to 70-80% confluency. Cultured cells (2 x 10^6^ cells/mL) were treated with non-stressed monomers and stressed aggregates of mAb 4713 at a final concentration of 2 μg/mL and incubated at 37 °C for 30 minutes. Unbound antibody mixtures were washed away and resuspended in FACS buffer. Then, dead cells were labelled with propidium iodide (PI) and incubated in dark for 2 minutes at 25 °C. Finally, the percentages of dead cells were investigated using flow cytometry and a haemocytometer.

Furthermore, we evaluated the neutralizing activity of ADA by measuring the binding inhibition of mAb 4713 to its target receptor on Raji cells. First, undiluted 10 μL of mouse heart antisera were mixed with non-stressed mAb 4713 at 2 μg/mL final concentration. Next, Raji cells (2 x 10^6^ cells/mL) were treated with the mAb 4713-antisera mixture, followed by an incubation at 37 °C for 30 minutes. As above, the mixture was washed, the dead cells were labelled with PI, and the percentages of dead cells were investigated using flow cytometry and haemocytometer.

### 2.7. Statistical analysis

Statistical analysis was performed using a two-tailed Student *t*-test with MS Excel (Microsoft, Redmond, WA, USA). The *p* value < 0.05 was considered statistically significant and asterisks (*) symbol denote statistical significance (* *p* < 0.05, \*\**p* < 0.01, *** *p* < 0.001, ns, not significant).

## 3. Results

### 3.1. Biophysical nature of stressed and non-stressed mAb 4713

We measured the sizes (hydrodynamic radius, *R*_h_) of mAb 4713 under stressed and non-stressed conditions by DLS and SLS. The DLS measurements showed that non-stressed proteins remained monomeric at both 25 °C and 37 °C (Fig. 1a and Supplementary Fig. S2a), with an *R*_h_ of 5.1 ± 0.4 nm. In contrast, heating, stirring, and freeze-thawing led to subvisible aggregate formation with *R*_h_ of 41.2 ± 3.1 nm, 265.2 ± 104.9 nm, and 512.9 ± 84.2 nm, respectively (Fig. 1a). SLS measurements corroborated the DLS results, showing low scattering intensity for non-stressed proteins and markedly higher intensities for all three types of stressed aggregates at both 25 °C and 37 °C (Fig. 1b, Supplementary Fig. S2b).

**Fig. 1:**
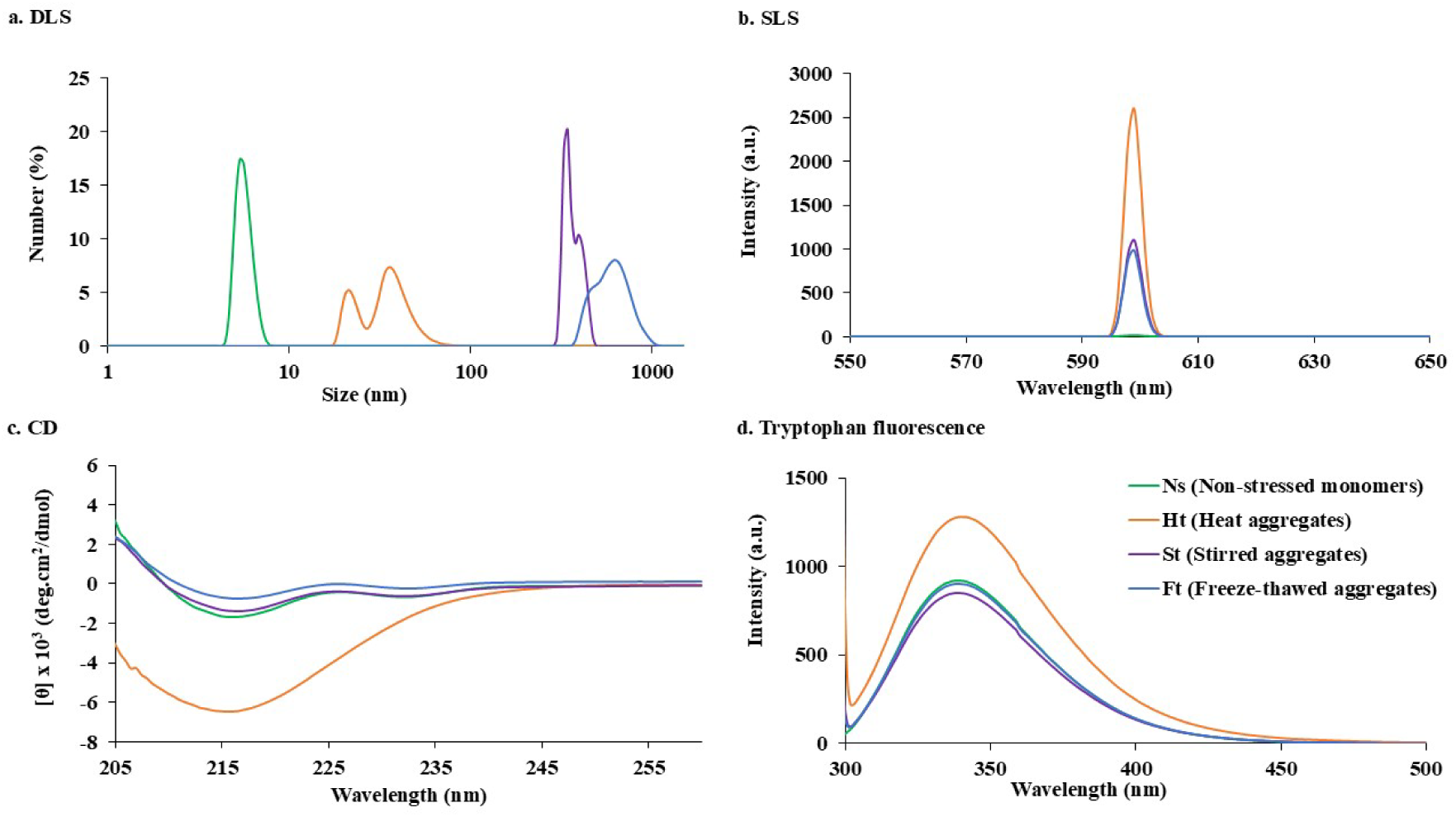
Biophysical properties of mAb 4713 and its aggregates. The **a.** DLS, **b.** SLS, **c.** far-UV CD, and **d.** tryptophan fluorescence spectra of stressed and non-stressed mAb 4713 measured at 25 °C. All measurements were done at 0.3 mg/mL protein concentration in 1x PBS buffer, pH 7.4. Line symbols are explained within the panels.

We analyzed the secondary structure contents and conformational state of the monomers and aggregates of mAb 4713 using far-UV CD, tryptophan fluorescence, and ANS fluorescence spectroscopy. The far-UV CD spectra of both stirred and freeze-thawed aggregates were identical to that of the monomeric protein, indicating retention of the native secondary structure at both 25 °C and 37 °C (Fig. 1c; Supplementary Fig. S2c; Supplementary Table S2). In contrast, heat-induced aggregates exhibited altered CD spectra, suggesting a loss of native secondary structure, attributable to mAb unfolding upon heating at 70 °C-consistent with previous observations [30]. Tryptophan and ANS fluorescence spectra of the monomers, and the stirred and freeze-thawed aggregates were also identical and typical of a natively and compactly folded protein at both 25 °C and 37 °C (Fig. 1d, 2a; Supplementary Fig. S2d, S3a). In contrast, the heat-induced aggregates showed altered tryptophan and ANS spectra with higher intensity, suggesting a non-native, loosely folded conformation, possibly related to a molten globule state [19,31,32].

Thermal stability assessed by TSA showed that the monomers, as well as the stirred and freeze-thawed aggregates, had similar melting curves (*T*_m_ ≈ 69.0 °C) (Fig. 2b; Supplementary Fig. S3b; Supplementary Table S3), supporting their native-like structures. In contrast, heat-induced aggregates displayed no cooperative melting curve, consistent with an unfolded state [24]. DSC analysis of the heat aggregates lacked a distinct thermal transition peak, indicating that incubation at 70 °C led to irreversible aggregate formation (Fig. 2c). On the other hand, the stirred and freeze-thawed aggregates were nearly as natively folded as the non-stressed monomers, as evidenced by their overlapping first endothermic peaks. However, the second endothermic peaks of the aggregates did not fully overlap, possibly reflecting minor structural alterations in the multidomain mAb 4713 due to the applied stresses [33]. Finally, limited proteolysis confirmed that the biochemical stabilities of the monomers and native-like aggregates were comparable, whereas that of the heat-induced, denatured aggregates was substantially reduced (Fig. 2d; Supplementary Fig. S3c) [19].

**Fig. 2:**
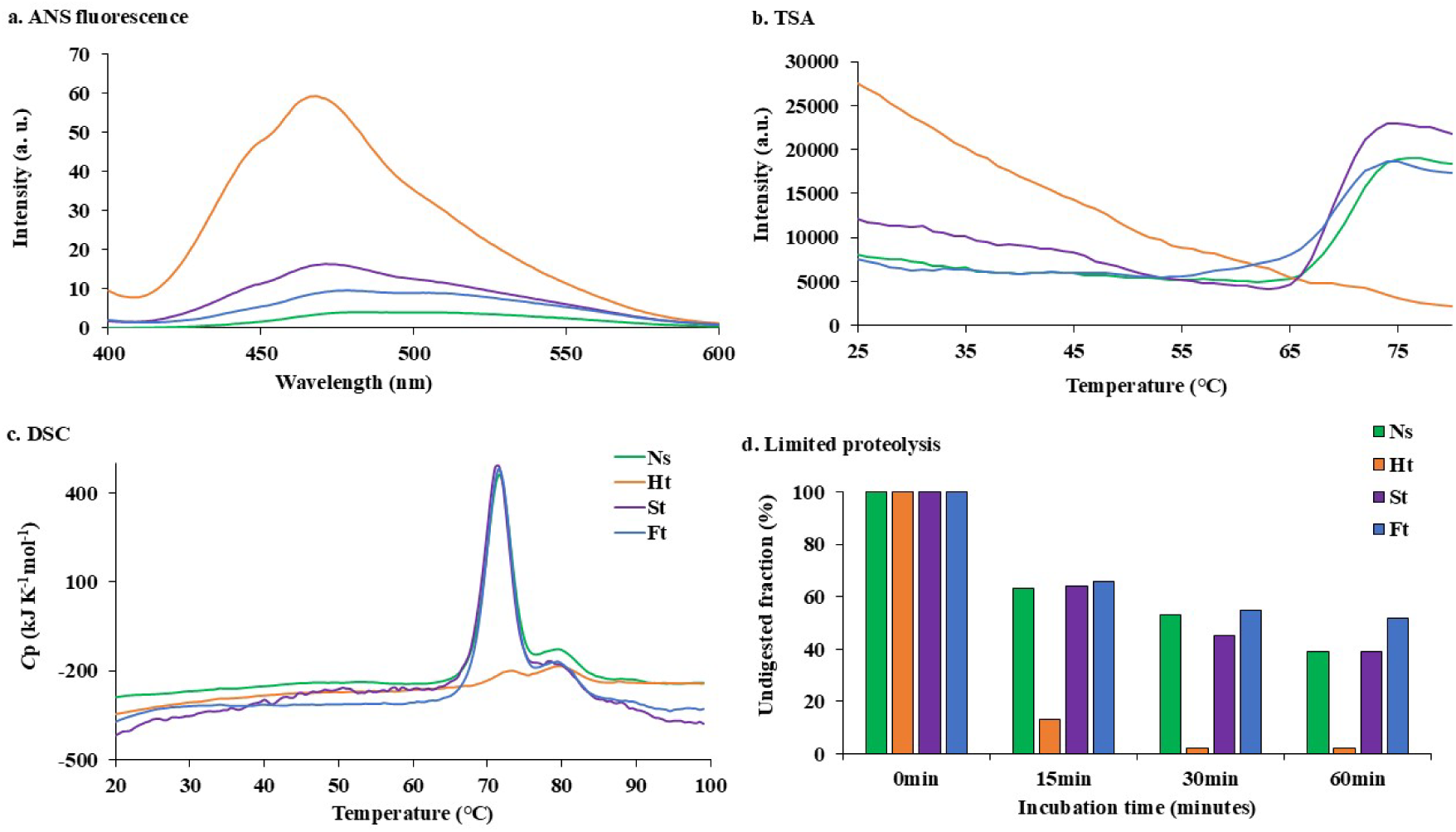
Biophysical properties of mAb 4713 and its aggregates. The **a.** ANS spectra, **b.** melting curves of the proteins assessed by TSA, **c.** DSC thermograms at the scan rate of +1.0 °C/min. All the measurements were done under the same conditions as mentioned in Fig. 1. **d.** To assess the biochemical stability, non-stressed and stressed mAb 4713 at 0.15 mg/mL concentration were treated with trypsin at 175 µg/mL final concentration and incubated at 37 °C for different time durations. The fraction of the proteins that remained intact after digestion was calculated from the SDS-PAGE image using CS analyzer 4 software (ATTO, Tokyo Japan) and undigested fractions are shown as percentages. The SDS gels image is shown in Supplementary Fig. S3c.

### 3.2. Anti-mAb 4713 immune responses in Jcl: ICR mice

We investigated the generation of anti-mAb 4713 antibodies (ADA) and the formation of memory T-cell populations using ELISA and flow cytometry, respectively, in the Jcl: ICR mouse model. The aggregates were freshly prepared before each immunization, and their sizes (*R*_h_) were confirmed by DLS (Supplementary Fig. S3d).

As expected, the non-stressed monomeric protein (Ns) was non-immunogenic even after the third dosing, as evidenced by the absence of both anti-mAb 4713 IgM and IgG antibody titers (Fig. 3a, 3b). The heat (Ht)-and stirred (St)-aggregates induced a modest and a strong immune response respectively, after the second dose, with IgM titers approximately 2-fold and 10-fold higher than those elicited by the monomeric protein (Fig. 3a; Supplementary Fig. S4a, S4b), consistent with previous reports [34]. In addition, a similar pattern was observed for the IgG response, with heat- and stirred-aggregates producing approximately 21-fold and 72-fold higher IgG titers, respectively, compared to the monomeric protein (Fig. 3b; Supplementary Fig. S4c, S4d). The decrease in IgM titers after the third dose, along with the subsequent increase in IgG titers, is indicative of Ig class switching [35]. On the other hand, freeze-thawed aggregates (Ft), despite being the largest in size (500-700 nm), were not immunogenic (Fig. 3b, Supplementary Fig. S4e). We further analysed IgG subclass to characterize the immune response against the aggregates. A high IgG1/IgG2a ratio indicated that stirred aggregates elicited a T_H_2-biased humoral immune response (Supplementary Fig. S4f) [36].

**Fig. 3:**
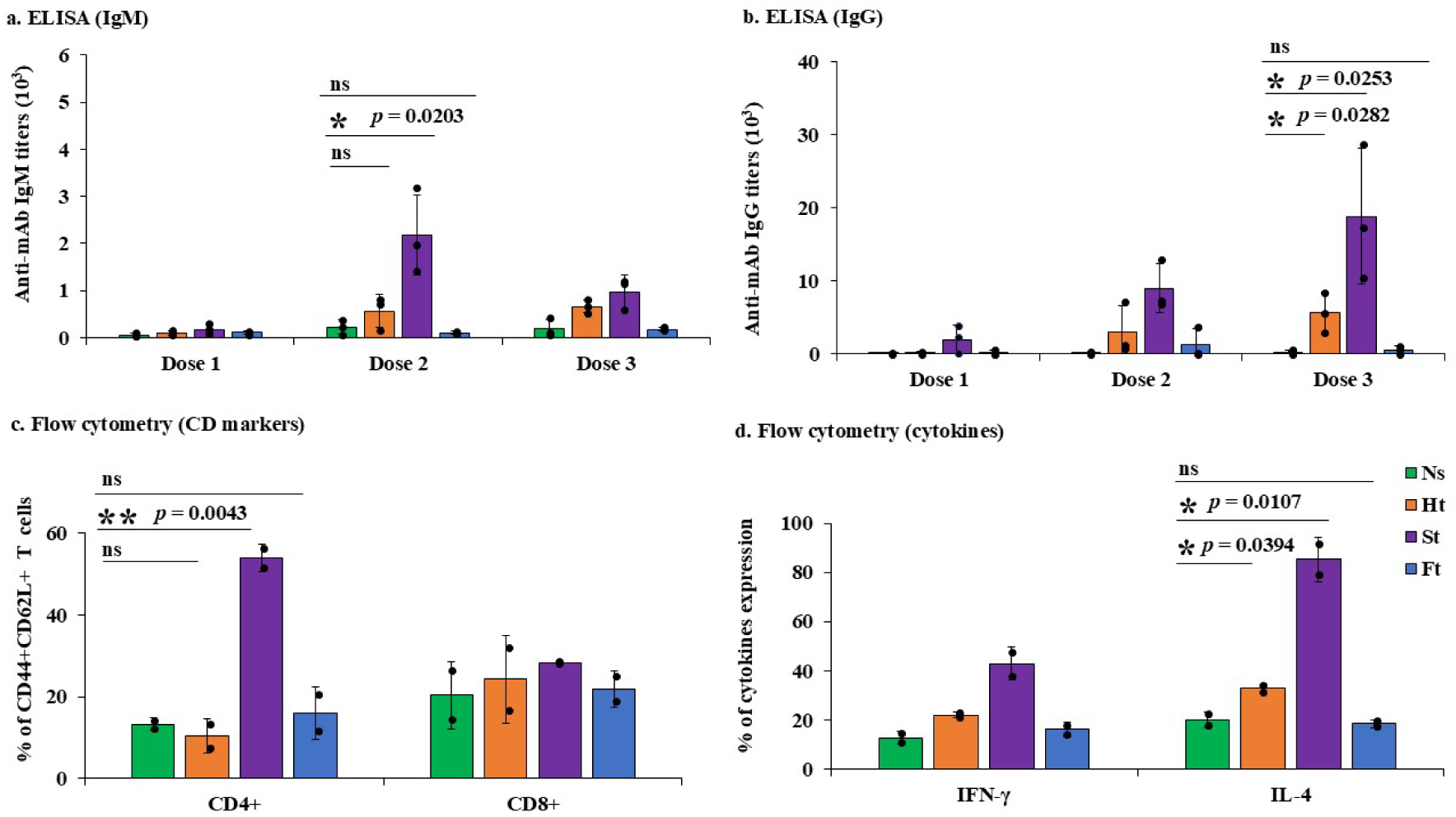
Anti-mAb 4713 antibody (ADA) responses in Jcl: ICR mouse model. Dose-specific anti-mAb 4713 **a.** IgM and **b.** IgG titers. Dose 1, 2, 3 represent the antibody titers using antisera from TB-2, 4, and 6 samples, respectively. ELISA plates were coated with non-stressed mAb 4713 at 100 ng/mL (IgM) and 40 ng/mL (IgG) concentrations. The average antibody titers of all the mice within a group (*n* = 3) are shown. **c.** Comparative CD44^+^-CD62L^+^ splenic T-lymphocytes (CD4^+^ and CD8^+^) and **d.** cytokines (IFN-γ and IL-4) expression pattern of CD4^+^ T-lymphocytes shown as percentage were assessed by flow cytometry. For flow cytometric analysis, two individual mice from each group were used (*n* = 2). Closed black circles (●) show the results for each mouse. Error bars represent the standard deviations. Statistical analysis was performed using a two-tailed Student *t*-test with MS Excel (Microsoft, Redmond, WA, USA). Here, the *p* value < 0.05 was considered statistically significant and asterisks (*) symbol denote statistical significance (* *p* < 0.05, \*\**p* < 0.01, *** *p* < 0.001, *p* > 0.05, ns, not significant). Ns, Ht, St, and Ft represent the mice groups that received mAb 4713 monomers, heat, stirred, and freeze-thawed aggregates, respectively.

Next, we investigated the expression of cell surface CD markers and cytokines in mouse splenic T lymphocytes by flow cytometry. Co-expression of CD44 and CD62L on both helper (CD4) and cytotoxic (CD8) T cells revealed that only the group immunized with stirred aggregates exhibited a markedly higher percentage of CD44 -CD62L helper T (CD4) cells, suggesting the formation of strong humoral immune memory (Fig. 3c; Supplementary Fig. S5) [37]. In addition, this group showed higher IL-4 expression relative to IFN-γ, further supporting a T_H_2-dependent humoral immune response (Fig. 3d) [36].

### 3.3. Therapeutic cytotoxicity and inhibition of the cytotoxicity by ADA

The mAb 4713 was developed as a therapeutic antibody targeting HLA class-II molecules for the treatment of several lymphomas, including Hodgkin, Burkitt, B-cell, NK-cell, and adult T-cell leukemia (ATL) [22]. We thus examined the therapeutic activity i.e. the anti-tumor cytotoxicity of mAb 4713 using Raji cells, a Burkitt lymphoma cell line expressing HLA class-II, via flow cytometry and direct cell counting by light microscopy, as previously described [22]. Cell death assay results revealed that the stirred and the freeze-thawed aggregates exhibited cytotoxicity comparable to that of the non-stressed monomeric mAb 4713 (25-60%), whereas heat aggregates showed no detectable cytotoxicity (only 8%, same as negative control (NC)), indicating a complete loss of therapeutic function (Fig. 4). Somewhat unexpectedly, these findings suggest that mAb 4713 can retain full therapeutic activity even when aggregated, provided that the structural integrity of mAb 4713 is preserved. In contrast, activity was lost when mAb 4713 was aggregated in a denatured state, as we anticipated.

**Fig. 4:**
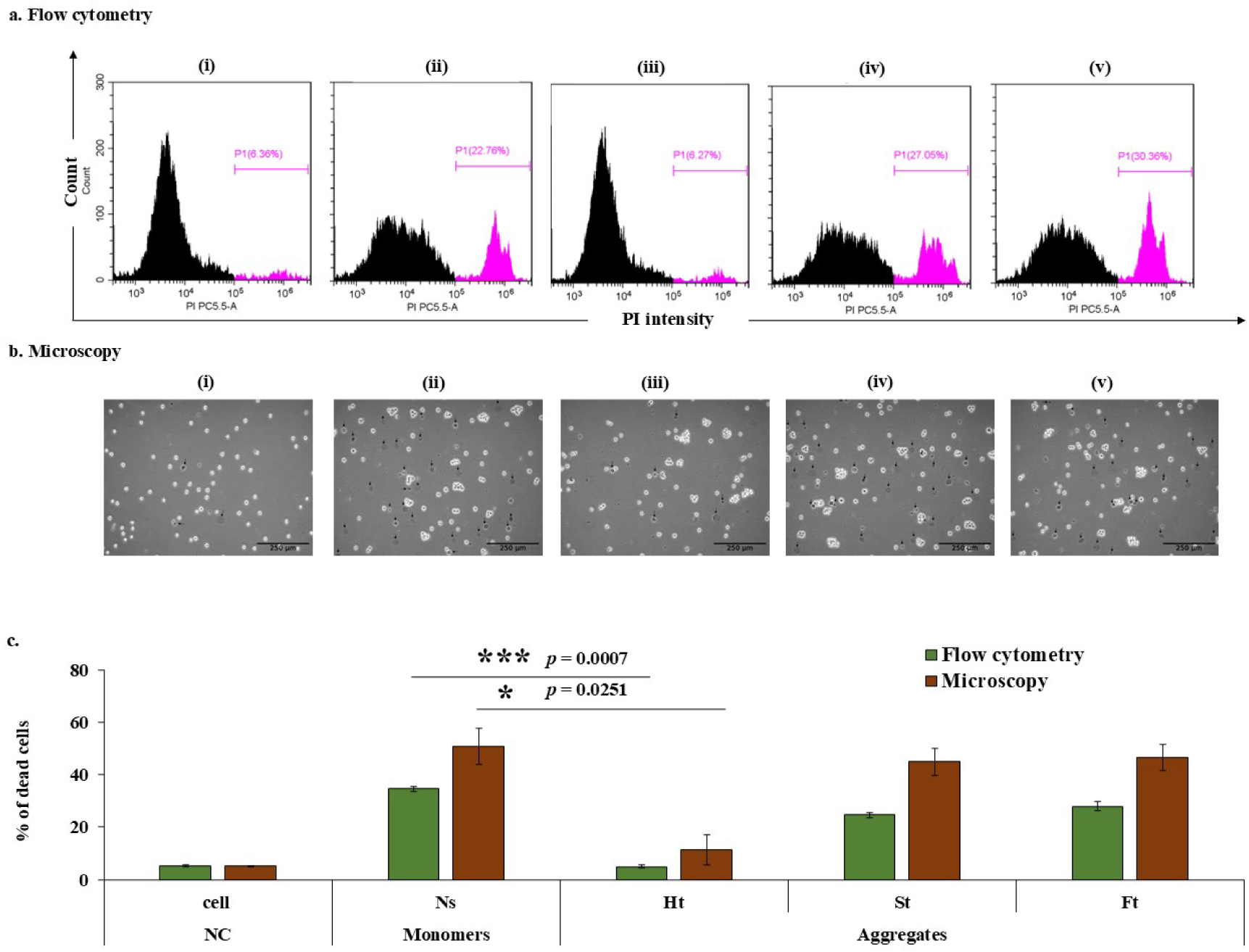
Anti-tumor cytotoxicity by mAb 4713 and its aggregates. The cytotoxicity of mAb 4713 monomers and aggregates analysed by incubating Raji cells at 37 °C in the absence (negative control, NC) and presence of mAb 4713 non-stressed monomers and stressed aggregates. **a.** The cytotoxicity analysed by flow cytometry by staining the dead cells (purple peak) with propidium iodide (PI). **b.** Light microscopy images showing trypan blue stained cells. Dead cells are indicated by black arrow marks. Scale bar: 250 µm. Here, , , , , and represent cell NC, Ns monomers, and Ht, St, and Ft aggregates respectively. **c.** The dead cells counted by both flow cytometry and microscopy shown as percentages. Two individual measurements were done and averaged. Error bars represent standard deviations. Statistical analysis was performed as mentioned in Fig. 3.

Next, to evaluate the functional consequences of ADA generation, we assessed whether antisera from immunized mice could inhibit the cytotoxicity of mAb 4713. Specifically, we found that heart antisera raised against stirred aggregates completely inhibited the cytotoxicity (only 7%), indicating the presence of neutralizing ADA induced by structurally intact aggregates (Fig. 5) [4,5,7]. In contrast, heart antisera from mice immunized with monomeric mAb 4713 or other aggregate types had no inhibitory effect on cell death (Fig. 5), consistent with their non-or mild-immunogenic profiles (Fig. 3).

**Fig. 5.**
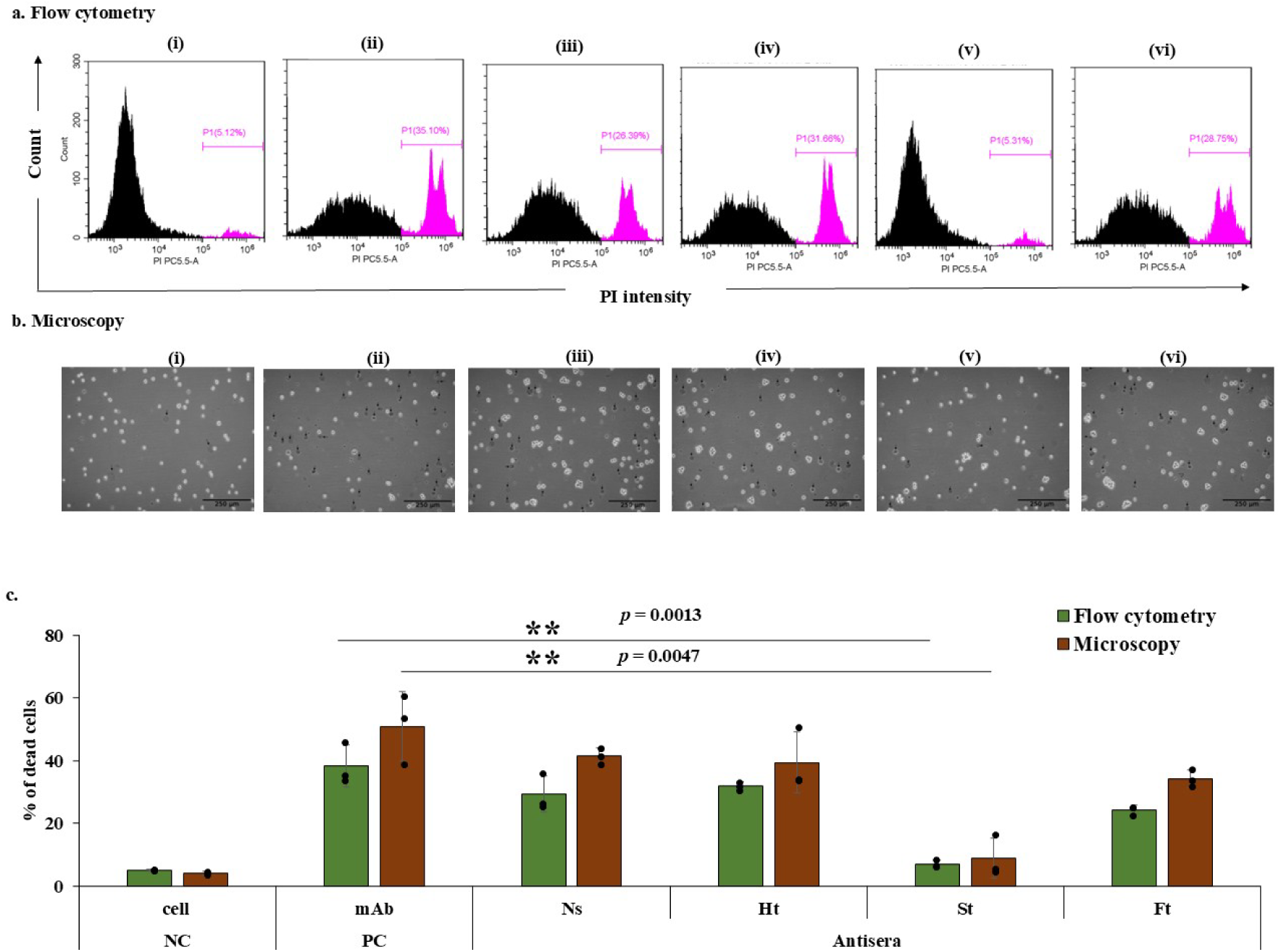
Inhibition of anti-tumor cytotoxicity by ADA. The inhibition of cytotoxicity analysed by incubating Raji cells at 37 °C in the absence (negative control, NC), presence (positive control, PC) of non-stressed mAb 4713, and mixture of antisera and mAb 4713. **a.** The cytotoxicity analysed by flow cytometry by staining the dead cells (purple peak) with propidium iodide (PI). **b.** Light microscopy images showing trypan blue stained cells. Dead cells are indicated by black arrow marks. Scale bar: 250 µm. Here, , , , , , and represent cell NC, mAb PC, and antisera from Ns, Ht, St and Ft aggregates injected mice groups, respectively. **c.** The dead cells counted by both flow cytometry and microscopy shown as percentages. The averages of all the mice within a group (*n* = 3) are shown. Closed black circles (●) show the results for each mouse. Error bars represent standard deviations. Statistical analysis was performed as mentioned in Fig. 3. Ns, Ht, St, and Ft represent the mice groups that received mAb 4713 monomers, heat, stirred, and freeze-thawed aggregates, respectively.

## 4. Discussion

Although aggregation remains a major concern in therapeutic antibody development due to its potential to induce neutralizing anti-drug antibodies (ADA) [3–5,7,8,10,12], the biophysical characteristics of protein aggregates and their impact on immune activation after administration remain under debate [16–20,38]. Clarifying this relationship has been difficult, partly because both protein aggregation and the resulting immune responses occur sporadically, making it challenging to establish a direct mechanistic link. To investigate the impact of the biophysical properties of therapeutic protein aggregates on unwanted immune responses, we employed a well-controlled system, both in terms of aggregate formation and biological evaluation. We generated three types of aggregates and assessed their impacts on immunogenicity *in vivo*.

Although all aggregates were subvisible (less than 1 micron), they elicited markedly different immune responses, as reflected by their IgM and IgG titers (Fig. 3). This indicates that aggregate size alone did not determine the immunogenicity of the proteins [16]. Instead, the biophysical properties of the aggregates strongly influenced both immune activation and tumoricidal activity. While heat-induced denatured aggregates triggered only a mild immune response (Fig. 3), they completely lost their therapeutic effect (Fig. 4), suggesting that structural perturbation of a therapeutic protein can compromise its biological activity.

Here, we would like to emphasize that the generation of antibodies does not necessarily equate to the production of neutralizing anti-drug antibodies (ADA) [4,5]. Non-neutralizing ADAs that do not interfere with drug activity are not immediately harmful, at least at the cellular level. Indeed, although heat-induced aggregates triggered a mild immune response (Fig. 3), the resulting antisera did not impair the function of mAb 4713 (Fig. 5), supporting the idea that non-neutralizing ADA did not compromise therapeutic efficacy [4,5,7]. Whether this lack of inhibition is due to the relatively low antibody titers or because the ADA, generated against denatured epitopes, are intrinsically unable to recognize the native, folded form of mAb 4713 remains an open question for future investigation. In contrast, the stirred aggregates retained their native structural integrity and function (Fig. 1, 2, 4), yet induced a robust immune response that ultimately inhibited the drug’s anti-tumor cytotoxicity (Fig. 3, 5). This can be interpreted as the emergence of neutralizing ADA [4,5,7].

These findings lead us to conclude that native-like aggregates can elicit a strong antibody response, consistent with previous reports showing that native-like aggregates of recombinant factor VIII are highly immunogenic [39]. Additionally, earlier studies have demonstrated that the immunogenicity of subunit vaccine candidates can be enhanced by forming native-like aggregates derived from small, non-immunogenic proteins through the use of solubility controlling peptide (SCP) tags [26,27].

Interestingly, freeze-thawed aggregates - despite being the largest (500-700 nm) and retaining both native structure and therapeutic function - did not elicit a detectable immune response (Fig. 3, Supplementary Fig. S4e). Their relatively low colloidal stability may help explain this lack of immunogenicity (Supplementary Fig. S6, S7). In contrast, the stirred aggregates, which were smaller (200-300 nm) but exhibited higher colloidal stability (Supplementary Fig. S6), were highly immunogenic (Fig. 3). Our *in vitro* analysis showed that the freeze-thawed aggregates gradually dissociated into monomers within 24 hours, confirming their lower colloidal stability (Supplementary Fig. S6, S7). Although all aggregates were freshly prepared just prior to immunization, it is possible that colloidally stable aggregates are more immunogenic due to their prolonged retention at the injection site, as previously suggested [16,40].

## 5. Conclusion

To the best of our knowledge, this is the first study to directly link the formation of distinct types of protein aggregates with the induction of neutralizing anti-drug antibodies (ADA). We emphasize that ADA formation is largely a sporadic phenomenon, which makes it difficult to characterize systematically. However, it can be conceptually stratified into at least three stages: (1) the formation of protein aggregates, (2) induction of an immune response, and (3) loss of therapeutic function.

In addition, by using well-controlled systems and detailed characterization at each of these stages, we demonstrated that therapeutic protein aggregates can induce neutralizing ADA *in vivo*, which completely abolished the therapeutic activity of the drug. Notably, not all types of aggregates induced neutralizing ADA; for example, heat-induced aggregates triggered a mild immune response but did not result in neutralizing antibodies. Altogether, our findings suggest that therapeutic protein drugs should be monitored not only for aggregate size— especially under external stresses—but also for their colloidal stability and biophysical properties, in order to preserve therapeutic efficacy from production to clinical use.

## Institutional review board statement

All mouse experiments were conducted under the Japanese animal experimentation law and the TUAT regulation, which the committee reviewed and approved on 25 March 2024 [R06-29].

## CRediT authorship contribution statement

**Sanjida Yesmin:** Writing – review & editing, Writing – original draft, Visualization, Methodology, Investigation, Formal analysis. **M. Monirul Islam:** Writing – review & editing, Supervision, Methodology, Investigation, Formal analysis. **Md. Din Islam:** Writing – review & editing, Methodology. **Hiromichi Tsurui:** Writing – review & editing, Supervision, Conceptualization. **Tomonori Saotome:** Methodology, Investigation. **Shuji Matsuoka:** Resources. **Yutaka Kuroda:** Writing – review & editing, Supervision, Funding acquisition, Conceptualization.

## Acknowledgments

We are grateful to Patricia McGahan for English proofreading, Professor Shun-ichi Kidokoro for discussion with DSC data, and all the members of Kuroda’s laboratory for advice and discussion.

## Data availability statement

All data generated or analysed during this study are included in this article and its supplementary information file.

## Abbreviations

ADA: anti-drug antibody
CD: circular dichroism
DLS: dynamic light scattering
DSC: differential scanning calorimetry
ELISA: Enzyme-linked immunosorbent assay
Ig: immunoglobulins
mAb: monoclonal antibody
PBS: phosphate-buffered saline
SLS: static light scattering
TSA: thermal shift assay

---

**Table 1:** Summary of the biophysical nature of mAb 4713 aggregates and their impacts on immune induction and therapeutic activity inhibition. The aggregate sizes (hydrodynamic radius, *R*h in nm), structural integrity, colloidal stability, anti-tumor cytotoxicity, anti-mAb IgG titers of heart antisera, and the percentages of cell death by neutralizing ADA are shown.

| Sample<sup>a</sup> | Size (nm)<sup>b</sup> | Type<sup>c</sup> | Colloidal stability<sup>d</sup> | Anti-tumor cytotoxicity<sup>e</sup> | IgG Titers<sup>f</sup> | Cell death by ADA<sup>g</sup> |
| --- | --- | --- | --- | --- | --- | --- |
| Ns | 4-6 | native | - | 34% | 138 | 29% |
| Ht | 30-50 | denatured | high | 5% | 6222 | 31% |
| St | 200-300 | native | high | 24% | 32014 | 7% |
| Ft | 500-700 | native | low | 27% | 769 | 24% |

<sup>a</sup>Ns, Ht, St, and Ft represent non-stressed monomers, heat, stirred, and freeze-thawed aggregates, respectively as mentioned in Fig. 1.
<sup>b</sup>Sizes of the aggregates ( $R_h$ , nm) were determined by DLS.
<sup>c</sup>Types of aggregates were assessed by CD, tryptophan spectroscopy, and TSA.
<sup>d</sup>Colloidal stability was monitored by incubating the aggregates at 37 °C.
<sup>e</sup>Anti-tumor cytotoxicity of the monomers and aggregates was assessed by cell death assay using flow cytometry.
<sup>f</sup>Anti-mAb IgG titers were assessed by ELISA, coating with non-stressed monomeric mAb 4713.
<sup>g</sup>Cell death by ADA was assessed by cell death assay using flow cytometry.

