## Supplementary_information for "“Biophysical and Colloidal Properties Govern Anti-Drug Antibody Generation: A Case Study Using the Tumoricidal mAb 4713”"

**Supplementary Table S1: Immunization scheme**

| **Group** | **Antigen** | **Antigen concentration**  **(μg/100μL/dose/mouse)** | **Buffer** | **Mice number** |
| --- | --- | --- | --- | --- |
| 1 | Ns | 30 | PBS, pH: 7.4 | 3 |
| 2 | Ht | 30 | PBS, pH: 7.4 | 3 |
| 3 | St | 30 | PBS, pH: 7.4 | 3 |
| 4 | Ft | 30 | PBS, pH: 7.4 | 3 |

**Supplementary Table S2: Secondary structure contents of mAb 4731 and its aggregates by BeStSel.** The secondary structure contents of non-stressed and stressed mAbs were evaluated by the BeStSel online tool using far-UV CD data shown in Figure 2A [1].

| **Sample** | **α helix (%)** | **Anti-parallel β sheets (%)** | **Parallel β sheets (%)** | **Turn** | **Others (%)** |
| --- | --- | --- | --- | --- | --- |
| Ns | 0.0 | 44.0 | 0.0 | 15.1 | 40.9 |
| Ht | 5.6 | 40.2 | 2.4 | 14.6 | 37.3 |
| St | 0.0 | 44.5 | 0.0 | 16.4 | 39.1 |
| Ft | 0.0 | 41.5 | 0.0 | 16.2 | 42.3 |

**Supplementary Table S3: Melting temperature of native monomers and aggregates.** The melting temperature of non-stressed and stressed aggregates evaluated by TSA.

| **Sample** | **Melting temperature, *T*m (° C)** |
| --- | --- |
| Ns | 69.8 |
| Ht | - |
| St | 69.2 |
| Ft | 69.7 |

**Supplementary Figures**

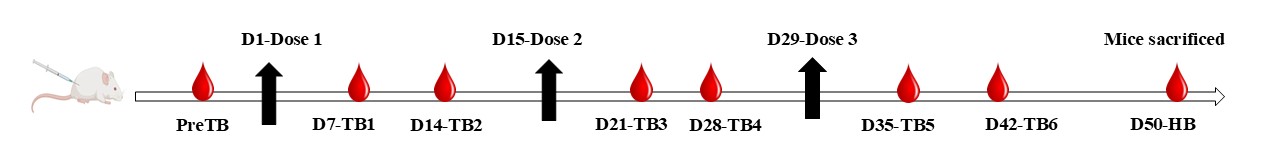

**Supplementary Fig. S1: Immunization schedule.** Mice immunization dosing schedule and sample collection days (D) are shown. Blood samples were collected from mice by tail bleeding (TB) and heart bleeding (HB).

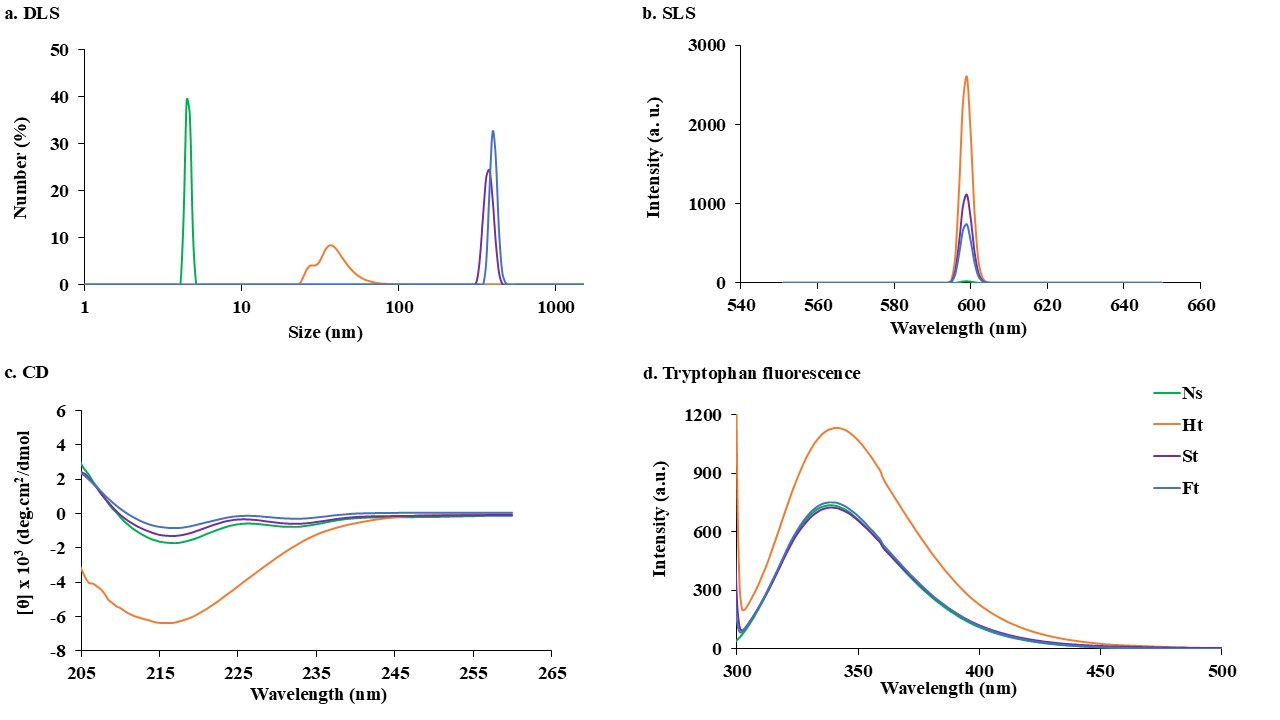

**Supplementary Fig. S2: Biophysical properties of mAb 4713 and its aggregates.** The **a.** DLS, **b.** SLS, **c.** far-UV CD, and **d.** tryptophan fluorescence spectra of stressed and non-stressed mAb 4713 measured at 37 ºC. Line symbols are explained within the panels.

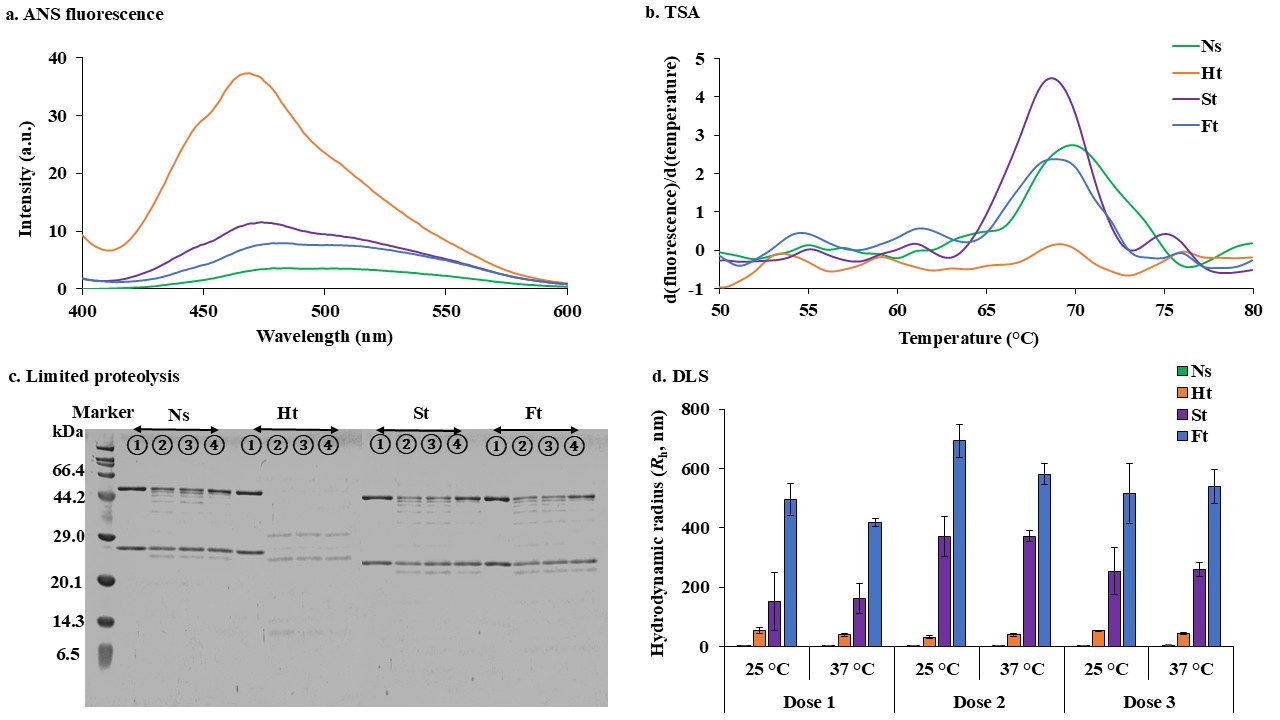

**Supplementary Fig. S3: Biophysical properties of mAb 4713 and its aggregates.** The **a.** ANS spectra of stressed and non-stressed mAb 4713 measured at 37 ºC, **b.** The derivative plot of the changes in fluorescence vs. temperature assessed by TSA, and **c.** SDS-PAGE gel for limited proteolysis. The biochemical stability of the mAb 4713 monomers and aggregates at 0.15 mg/mL concentration were evaluated treating with trypsin and the fraction of the sample was collected at different time durations and analysed by SDS-PAGE. Circled 1, 2, 3, 4 indicate the 0-, 15-, 30- and 60- minutes incubation sample bands respectively. **d.** The hydrodynamic radius (*R*_h_) of mAb 4713 monomers and aggregates were measured by DLS at 25 and 37 °C just before each immunization.

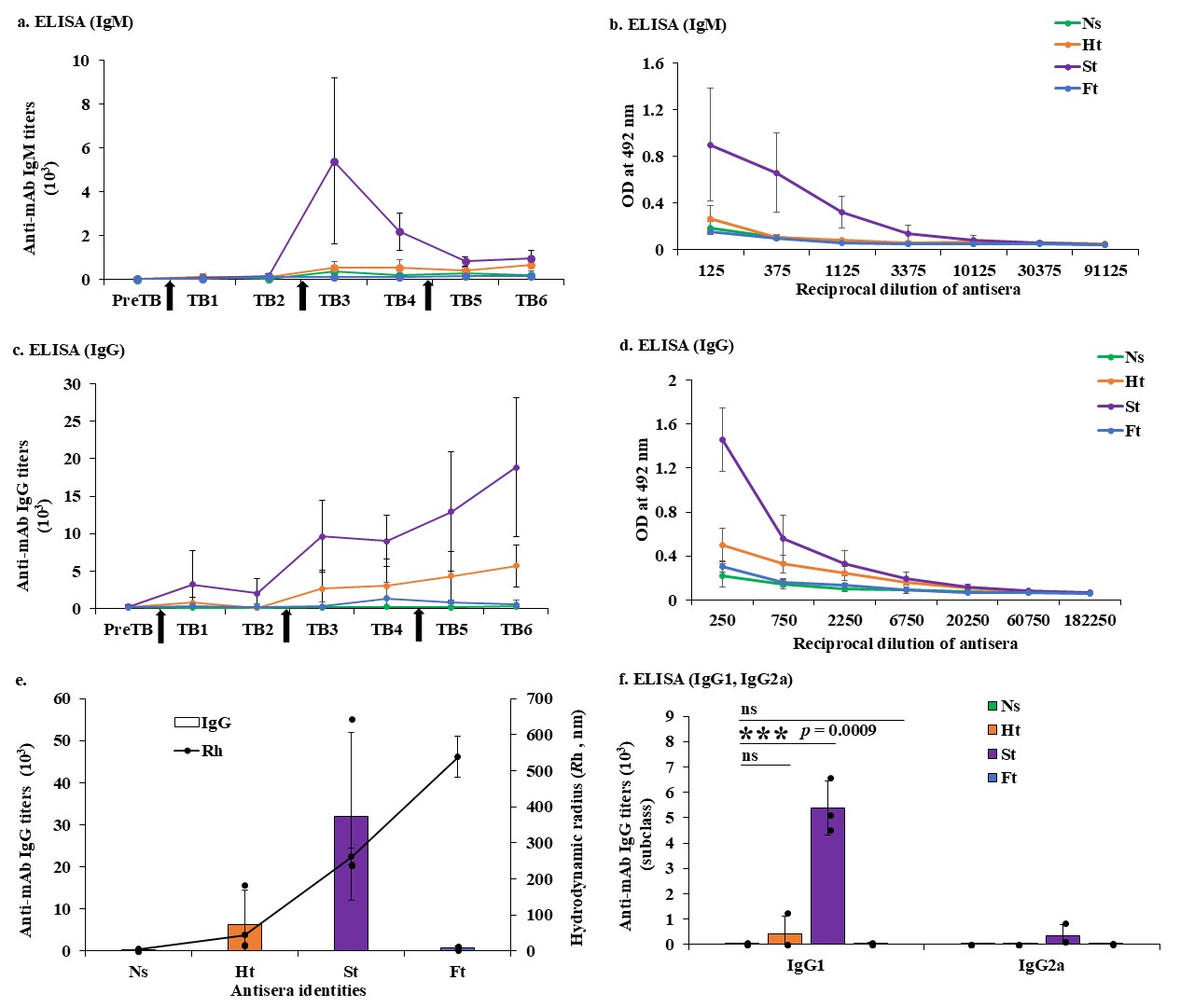

**Supplementary Fig. S4: Anti-mAb 4713 antibody (ADA) responses in Jcl: ICR mouse model.** **a.** Dose-dependent anti-mAb 4713 IgM titers, and **b.** OD at 492 nm of anti-mAb 4713 sera measured by [ELISA](https://www.sciencedirect.com/topics/immunology-and-microbiology/elisa) of the TB-3 samples for IgM response. **c.** Dose-dependent anti-mAb 4713 IgG titers and **d.** OD at 492 nm of anti-mAb 4713 sera measured by [ELISA](https://www.sciencedirect.com/topics/immunology-and-microbiology/elisa) of the TB-6 samples for IgG response. Black arrows and TB represent the immunization and antisera from tail-bleeding samples. **e.** Correlation between the antibody titers of heart antisera (primary axis) and aggregates hydrodynamic radius (*R*_h_) at 37 °C (secondary axis). **f.** IgG subclass (IgG1, IgG2a) determined by ELISA. ELISA plates were coated with non-stressed mAb 4713 at 100 ng/mL (IgM), 40 ng/mL (IgG) and 10 ng/mL (IgG1, IgG2a) concentrations. The average antibody titers of all the mice within a group (*n* = 3). Error bars represent standard deviations. Ns, Ht, St, and Ft represent the mice groups that received mAb 4713 non-stressed monomers, heat, stirred, and freeze-thawed aggregates, respectively.

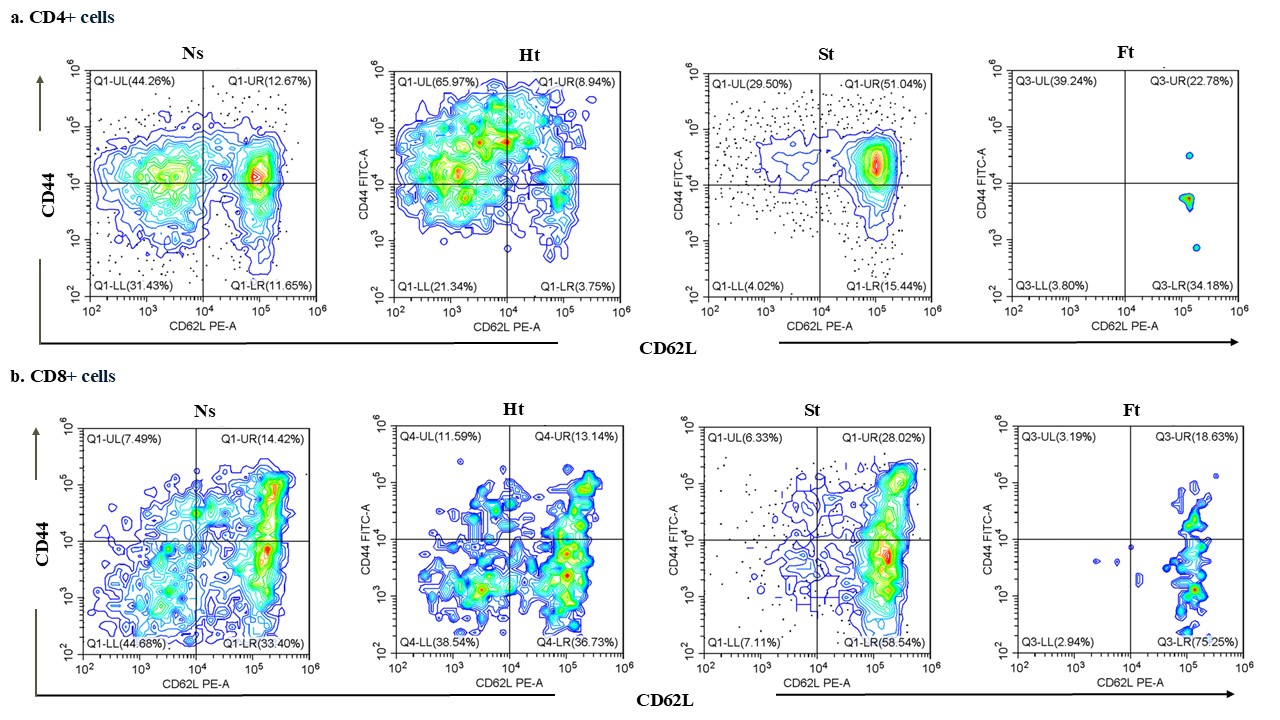

**Supplementary Fig. S5: Flow cytometry analysis of cell surface CD markers.** Dot plots of the differential expression of CD44 and CD62L on **a.**CD4^+^ and **b.**CD8^+^ cells of one mouse from each group. Ns, Ht, St, and Ft represent the mice groups that received mAb 4713 non-stressed monomers, heat, stirred, and freeze-thawed aggregates, respectively.

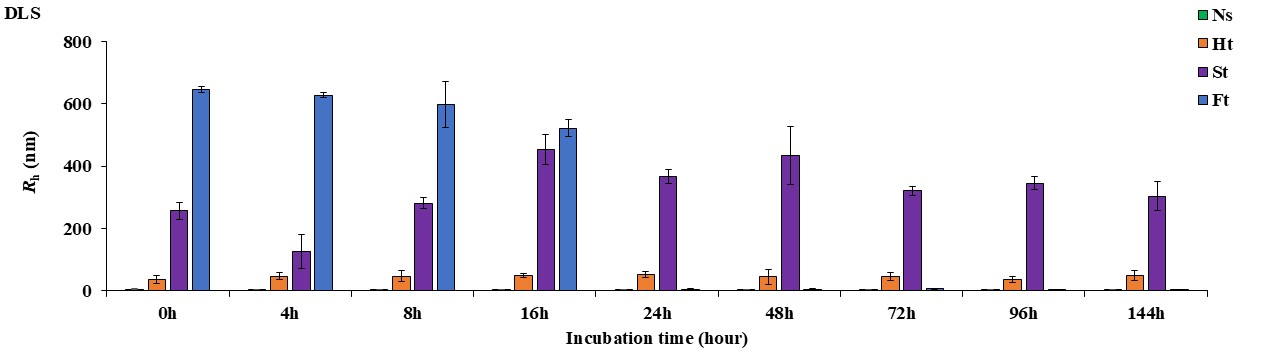

**Supplementary Fig. S6: Colloidal stability of mAb 4713 and its aggregates.** The non-stressed monomers and stressed mAb 4713 aggregates were incubated at 37 °C for different time durations and hydrodynamic radius (*R*_h_) measured at 37 °C by DLS. The hydrodynamic radius (*R*_h_) was computed from DLS’s number spectra.

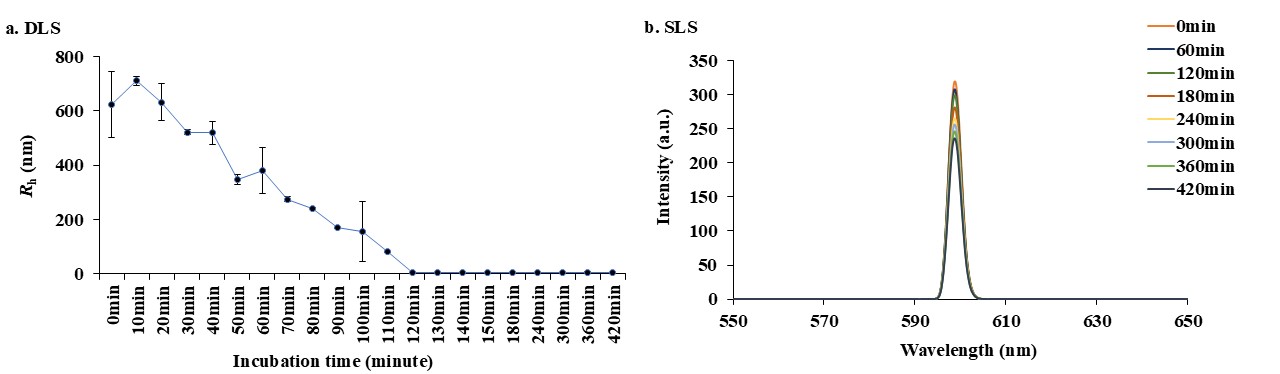

**Supplementary Fig. S7: Colloidal stability of freeze-thaw aggregates after 16 hours.** Freeze-thawed aggregates were incubated after 16 hours for different time durations at 37 °C and colloidal stability was determined by measuring hydrodynamic radius (*R*_h_) by DLS and static light intensity by SLS. The **a.** hydrodynamic radius (*R*_h_) and **b.** static light intensity are shown.

**Supplementary information references**

[1]	A. Micsonai, F. Wien, É. Bulyáki, J. Kun, É. Moussong, Y.-H. Lee, Y. Goto, M. Réfrégiers, J. Kardos, BeStSel: a web server for accurate protein secondary structure prediction and fold recognition from the circular dichroism spectra, Nucleic Acids Research 46 (2018) W315–W322. https://doi.org/10.1093/nar/gky497.
